# The substitutions L50F, E166A and L167F in SARS-CoV-2 3CLpro are selected by a protease inhibitor *in vitro* and confer resistance to nirmatrelvir

**DOI:** 10.1101/2022.06.07.495116

**Authors:** Dirk Jochmans, Cheng Liu, Kim Donckers, Antitsa Stoycheva, Sandro Boland, Sarah K Stevens, Chloe De Vita, Bert Vanmechelen, Piet Maes, Bettina Trüeb, Nadine Ebert, Volker Thiel, Steven De Jonghe, Laura Vangeel, Dorothée Bardiot, Andreas Jekle, Lawrence M Blatt, Leonid Beigelman, Julian A Symons, Pierre Raboisson, Patrick Chaltin, Arnaud Marchand, Johan Neyts, Jerome Deval, Koen Vandyck

**Affiliations:** KU Leuven, Department of Microbiology, Immunology & Transplantation, Rega Institute, Laboratory of Virology & Chemotherapy, Herestraat 49, 3000 Leuven, Belgium; Aligos Therapeutics, Inc., 1 Corporate Dr., 2nd Floor, South San Francisco, CA, USA; CISTIM Leuven vzw, Gaston Geenslaan 2, 3001 Leuven, Belgium; KU Leuven, Department of Microbiology, Immunology & Transplantation, Rega Institute, Laboratory of Clinical & Epidemiological Virology, Herestraat 49, Leuven, 3000, Belgium; Institute of Virology and Immunology, University of Bern, 3012, Bern, Switzerland; Department of Infectious Diseases and Pathobiology, Vetsuisse Faculty, University of Bern, Bern, Switzerland; Aligos Belgium BV, Gaston Geenslaan 1, 3001 Leuven, Belgium; Centre for Drug Design and Discovery (CD3), KU Leuven, Gaston Geenslaan 2, 3001 Leuven, Belgium; CISTIM Leuven vzw, Gaston Geenslaan 2, 3001 Leuven, Belgium; KU Leuven, Department of Microbiology, Immunology & Transplantation, Rega Institute, Laboratory of Virology & Chemotherapy, Herestraat 49, 3000 Leuven, Belgium and Global Virus Network (GVN)

## Abstract

The SARS-CoV-2 main protease (3CLpro) has an indispensable role in the viral life cycle and is a therapeutic target for the treatment of COVID-19. The potential of 3CLpro-inhibitors to select for drug-resistant variants needs to be established. Therefore, SARS-CoV-2 was passaged *in vitro* in the presence of increasing concentrations of ALG-097161, a probe compound designed in the context of a 3CLpro drug discovery program. We identified a combination of amino acid substitutions in 3CLpro (L50F E166A L167F) that is associated with > 20x increase in EC_50_ values for ALG-097161, nirmatrelvir (PF-07321332) and PF-00835231. While two of the single substitutions (E166A and L167F) provide low-level resistance to the inhibitors in a biochemical assay, the triple mutant results in the highest levels of resistance (6x to 72x). All substitutions are associated with a significant loss of enzymatic 3CLpro activity, suggesting a reduction in viral fitness. Structural biology analysis indicates that the different substitutions reduce the number of inhibitor/enzyme interactions while the binding of the substrate is maintained. These observations will be important for the interpretation of resistance development to 3CLpro inhibitors in the clinical setting.

**Abstract Importance:** Paxlovid is the first oral antiviral approved for treatment of SARS-CoV-2 infection. Antiviral treatments are often associated with the development of drug resistant viruses. In order to guide the use of novel antivirals it is essential to understand the risk of resistance development and to characterize the associated changes in the viral genes and proteins. In this work, we describe for the first time a pathway that allows SARS-CoV-2 to develop resistance against Paxlovid *in vitro*. The characteristics of *in vitro* antiviral resistance development may be predictive for the clinical situation. Therefore, our work will be important for the management of COVID-19 with Paxlovid and next generation SARS-CoV-2 3CLpro inhibitors.

## Introduction

There is an urgent need for potent and safe antiviral drugs for the treatment and prophylaxis of SARS-CoV-2 infections. Highly efficacious and safe viral protease inhibitors have contributed significantly to the effective treatment of infections with HIV and HCV. Coronaviruses have two proteases, the main protease 3CLpro (or Mpro) and the papain-like protease. 3CLpro is a cysteine protease that cleaves the two polyproteins (pp1a and pp1ab) of SARS-CoV-2 at eleven different sites, resulting in various non-structural proteins, which are key for viral replication [1]. The substrate of 3CLpro presents a distinct glutamine at the P1 site (Leu-Gln/Ser, Ala, Gly), while no known human proteases recognize this cleavage site [2, 3]. 3CLpro can thus be considered a highly attractive drug target for the development of SARS-CoV-2 antivirals [4]. The potential of 3CLpro inhibitors has become apparent with the development of nirmatrelvir (PF-07321332), a peptidomimetic reversible covalent inhibitor that is co-formulated with the pharmacokinetic enhancer ritonavir (the resulting combination being marketed as Paxlovid) [5]. When treatment is initiated during the first days after symptom onset, it results in substantial clinical benefit [6-9]. We recently demonstrated that nirmatrelvir is equipotent *in vitro* against the current SARS-CoV-2 variants of concern (VoC). Nirmatrelvir protects Syrian Golden hamsters from intranasal infection with different VoCs and prevents transmission to untreated co-housed sentinels [10, 11]. Other clinical candidate 3CLpro inhibitors include ensitrelvir (S-217622), a non-peptidic, non-covalent SARS-CoV-2 3CLpro inhibitor[12, 13], and PBI-0451 [14]. The clinical development of lufotrelvir, an intravenous pro-drug of PF-00835321 has been discontinued [15, 16].

Treatment with antivirals can result in the selection of resistant viral variants and subsequent therapeutic failure. This has been described extensively in the treatment of (chronic/persistent or acute) viral infections caused by HIV, HBV, HCV, herpesviruses or influenza [17, 18]. Importantly, transmission of resistant viruses has been reported for HIV and influenza [19, 20]. For SARS-CoV-2, selection of resistant isolates has only been described for remdesivir, a polymerase inhibitor. *In vitro* selection with remdesivir results in the emergence of resistance-associated mutations. Yet, in the clinical setting, treatment with remdesivir so far only led to the selection of mutations that are associated with low level resistance [21-23]. A causal effect between SARS-CoV-2 resistance to any replicase inhibitors and therapy failure has not yet been demonstrated, most likely because these inhibitors are not yet widely used in the clinic and/or resistant variants might have a fitness disadvantage.

Here we report on a pathway by which SARS-CoV-2 achieves significant resistance and cross-resistance to 3CLpro inhibitors during serial passage in cell culture in the presence of a first generation 3CLpro inhibitor, ALG-097161. This molecule was prepared as a tool compound in the context of a drug discovery program.

## Results

### SARS-CoV-2 acquires resistance to 3CLpro Inhibitors during passage with ALG-097161

ALG-097161 is a SARS-CoV-2 inhibitor with an EC_50_ of 0.59 μM when tested for inhibition of SARS-CoV-2 isolate GHB-0302, a prototypic Wuhan isolated from a Belgian patient, in VeroE6 cells (Table 1). It has no effect on the viability of uninfected host cells at concentrations up to 10 μM. Since VeroE6 cells have a high efflux of some chemotypes, all selection experiments, antiviral assays and toxicity assays on these cells were performed in the presence of the P-glycoprotein (Pgp) efflux inhibitor CP-100356 (0.5 μM) [24]. The antiviral activity of ALG-097161 can be ascribed to inhibition of 3CLpro (IC_50_ = 0.014 μM - Table 4) and there is no relevant inhibitory effect on human Cathepsin L (IC_50_ > 10 μM – Supplemental S1). For comparative reasons, the chemical structures of ALG-097161 and the clinical 3CLpro inhibitors nirmatrelvir, PF-00835231 and ensitrelvir are shown in Figure 1.

**Figure 1:**
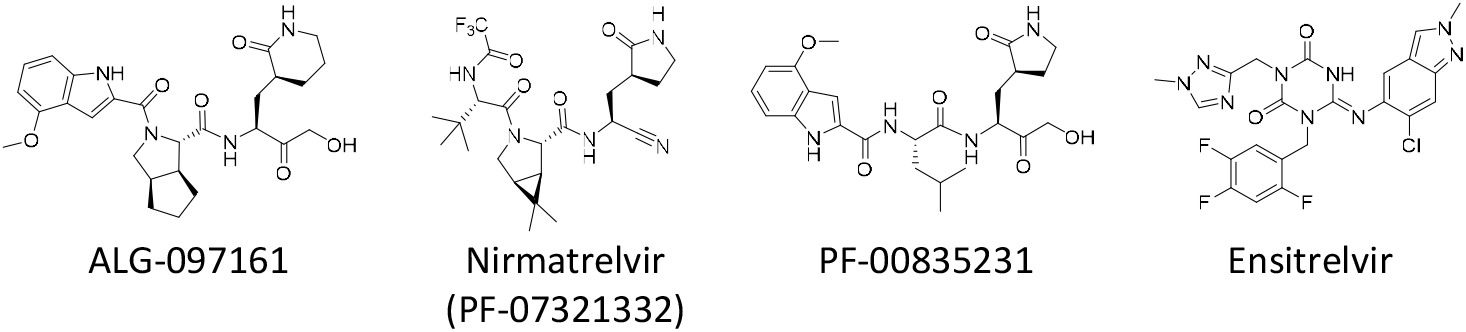
Chemical structures of different 3CLpro inhibitors.

**Table 1:** Phenotypic resistance associated with the L50F E166A L167F mutation profile as determined in VeroE6 cells.

|  | toxicity<br>VeroE6<br>CC<sub>50</sub> (μM) | SARS-CoV-2<br>WT-GHB-03021<br>EC<sub>50</sub> (μM) | SARS-CoV-2<br>L50F E166A L167F<br>EC<sub>50</sub> (μM) [FC]*** |
| --- | --- | --- | --- |
| **ALG-097161** | > 10<br>n=4 | 0.59*<br>(0.47 - 0.83)** n = 6 | 39 [63]<br>(13 - 47) n = 6 |
| **Nirmatrelvir** | > 10<br>n=4 | 0.12<br>(0.094 - 0.18) n = 8 | 6.1 [51]<br>(5.8 - 8.1) n = 5 |
| **PF-00835231** | > 10<br>n=4 | 0.19<br>(0.18 - 0.20) n = 2 | 4.4 [23]<br>(1.0 - 7.8) n = 5 |
| **Remdesivir**<br>**(GS-441524)\*\*\*\*** | > 10<br>n=4 | 0.79<br>(0.67 - 1.1) n = 8 | 1.7 [2.1]<br>(0.59 - 1.9) n = 6 |

\* Median value; \*\* 25th – 75th percentile, EC<sub>50</sub> = 50% effective concentration, \*\*\*FC = Fold Change of EC<sub>50</sub>,
\*\*\*\* GS-441524 is the parent nucleoside of remdesivir, it is intracellularly converted to the same active metabolite EC<sub>50</sub> (50% effective concentration), CC<sub>50</sub> (50% cytostatic concentration)
note: all assays on VeroE6 cells are performed in the presence of 0.5 μM CP-100356

To identify resistance development to 3CLpro inhibitors, we passaged SARS-CoV-2-GHB-0302 in VeroE6 cells in the presence of increasing concentrations of ALG-097161. The selection started at a concentration of 0.4 μM and for each next passage the virus was cultured at the same concentration, a 3x higher concentration and a 3x lower concentration. The culture demonstrating virus breakthrough, as observed by a significant cytopathic effect, at the highest concentration was then used for the next passage. We started this experiment in triplicate but for two trials we could not significantly increase the ALG-097161 concentration above 2 μM by passage 8. These cultures were therefore not further analyzed. In one culture however, the concentration of ALG-097161 could be increased gradually to 5 μM at passage 8 (p8, day 28) (Figure 2) and maintained for one passage at 15 μM, but that concentration had to be decreased again to 5 μM for the subsequent passages to allow viral replication until p12 (day 39) (Figure 2). Whole genome sequencing (Illumina) was performed on RNA purified from harvested supernatant at passages 5, 8 and 12. At p5, no mutations were identified in the 3CLpro gene but at p8 and p12 dominant mutations (>80% of the reads) were present that result in amino acid substitutions L50F E166A and L50F E166A L167F, respectively (Figure 2; Supplemental S2). As a control, SARS-CoV-2 was also passaged, at the same frequency, in the absence of compound and no mutations were identified in the 3CLpro gene at any passage (data not shown).

**Figure 2:**
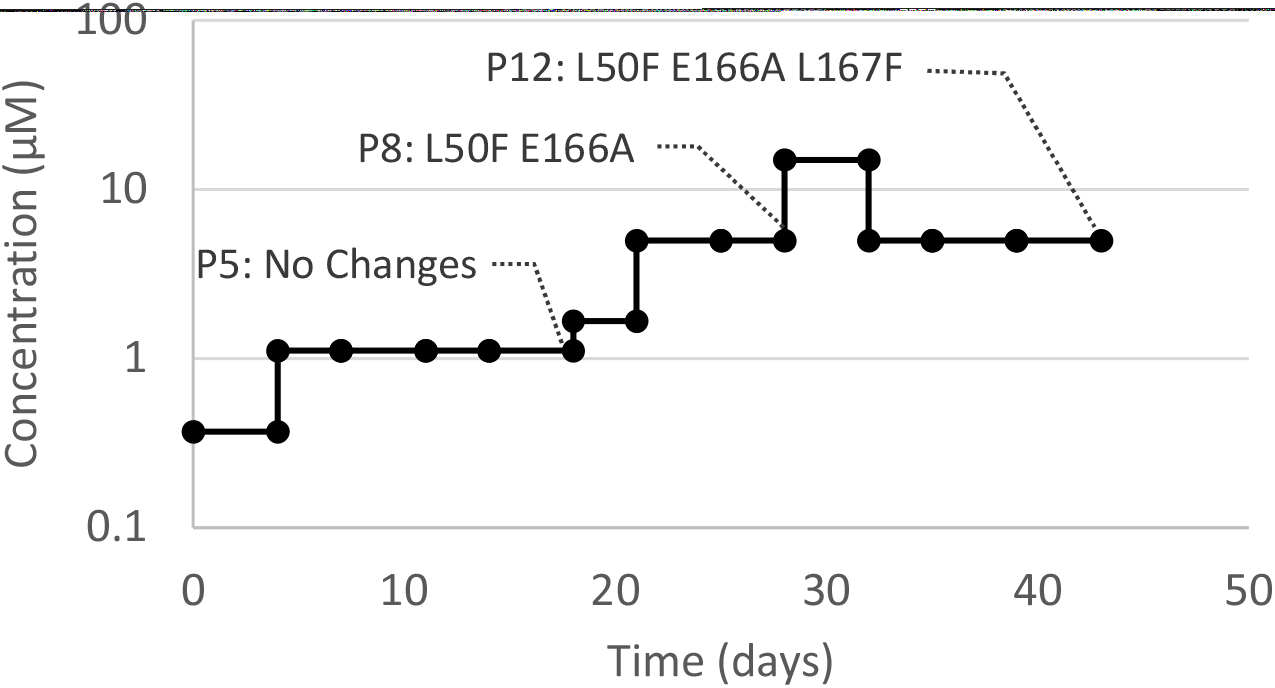
Passaging SARS-CoV-2-GHB (Wuhan) in VeroE6 cells in the presence of increasing concentrations of ALG-097161 (and the efflux inhibitor CP-100356). Selection was initiated at 0.4 μM. At every passage, several new cultures were started with the same concentration as well as a lower and a higher concentration. The passage with the highest compound concentration that could be maintained was selected for the following passage. At passage 5, 8 and 12, vRNA in the cell culture medium was sequenced. Substitutions in the 3CLpro that were found at these passages are indicated.

Analysis of the full genome of the selected viruses shows that also other nonsynonymous mutations became fixed (Supplemental S2). This can be explained by the fact that the nucleotide changes, resulting in the 3CLpro substitutions, occurred in a minor variant that also had these background mutations. Importantly, all these other changes are outside of the 3CLpro gene or the 3CLpro cleavage sites encoding sequences (Supplemental S2).

For phenotypic analysis, a new virus stock (p13) was grown in the presence of 4 μM ALG-097161, and the presence of L50F E166A L167F was confirmed by genotypic analysis. Antiviral testing with this stock showed strong resistance to ALG-097161 and cross-resistance to both nirmatrelvir and PF-00835231 (Table 1). The EC_50_ values for all these 3CLpro inhibitors are increased > 10x. As expected, the sensitivity for the polymerase inhibitor remdesivir remains unchanged.

In a second series of experiments, we investigated whether the observed amino acid substitutions in 3CLpro are sufficient for the resistance phenotype. To this end, we engineered, using an infectious clone [25], virus stocks with L50F, E166A L167F or L50F E166A L167F. Unfortunately, we were not able to generate virus with E166A or L167F alone. Genotypic analysis confirmed the presence of the mutations and revealed that there were no other nonsynonymous mutations (data not shown). Antiviral testing with these engineered viruses shows that L50F by itself is not associated with resistance while E166A L167F and L50F E166A L167F cause a 4.6x and 10.3x increase in EC_50_ for ALG-097161, respectively (Table 2). Again, cross-resistance is detected for nirmatrelvir (EC_50_ increased 29x with the L50F E166A L167F virus). At this stage of our study, ensitrelvir became available; we found that also this molecule shows cross-resistance (EC_50_ increased 44x with L50F E166A L167F). For all protease inhibitors tested we observe less resistance with the E166A L167F virus as compared with the L50F E166A L167F virus. Importantly, all the viruses analyzed remained fully sensitive to remdesivir.

**Table 2:** Phenotypic resistance associated with the L50F, E166A L167F and L50F E166A L167F mutation profile as determined by using reversed engineered SARS-CoV-2 viruses in the USA-WA1 background on VeroE6 cells.

|  | SARS-COV-2<br>WT-USA-WA1<br>EC<sub>50</sub> (μM) | EC<sub>50</sub> (μM) [FC]*** |  |  |
| --- | --- | --- | --- | --- |
|  |  | L50F | E166A L167V | L50F E166A L167V |
| **ALG-97161** | 0.60*<br>(0.51-0.71)** n=4 | 0.98 [1.6]<br>(0.83-1.1) n=4 | 2.9 [4.6]<br>(1.5-4.5) n=4 | 6.2 [10.3]<br>(4.8-7.3) n=4 |
| **Nirmatrelvir** | 0.11<br>(0.09-0.12) n=4 | 0.16 [1.5]<br>(0.12-0.2) n=4 | 1.1 [10.0]<br>(0.67-1.8) n=4 | 3.2 [29]<br>(2.3-3.5) n=6 |
| **Ensitrelvir** | 0.21<br>(0.18-0.27) n=4 | 0.24 [1.1]<br>(0.18-0.31) n=4 | 8.0 [38]<br>(2.8-12.7) n=4 | 9.3 [44]<br>(3.5-15) n=6 |
| **Remdesivir**<br>**(GS-441524)\*\*\*\*** | 0.86<br>(0.53-1.1) n=4 | 1.1 [1.3]<br>(0.8-1.1) n=4 | 1.5 [1.7]<br>(1.1-1.9) n=4 | 1.2 [1.4]<br>(0.91-1.7) n=6 |

\* Median value; \*\* 25th – 75th percentile, EC<sub>50</sub> = 50% effective concentration, \*\*\*FC = Fold Change of EC<sub>50</sub>,
\*\*\*\* GS-441524 is the parent nucleoside of remdesivir, it is intracellularly converted to the same active metabolite EC<sub>50</sub> (50% effective concentration)
note: all assays on VeroE6 cells are performed in the presence of 0.5 μM CP-100356

### Resistance to 3CLpro inhibitors in a cell-based reporter assay

The resistance profile was next confirmed in a previously described cell-based reporter assay of SARS-CoV-2 3CLpro enzymatic function [26]. In this gain-of-signal assay, inhibition of 3CLpro results in an increased eGFP signal. Introducing the three amino acid changes L50F E166A L167F into the construct resulted in a 23x and 28x loss of potency for ALG-097161 and nirmatrelvir respectively (Table 3). The resistance level of the triple mutant against PF-00835231 was 6.9x.

**Table 3:**
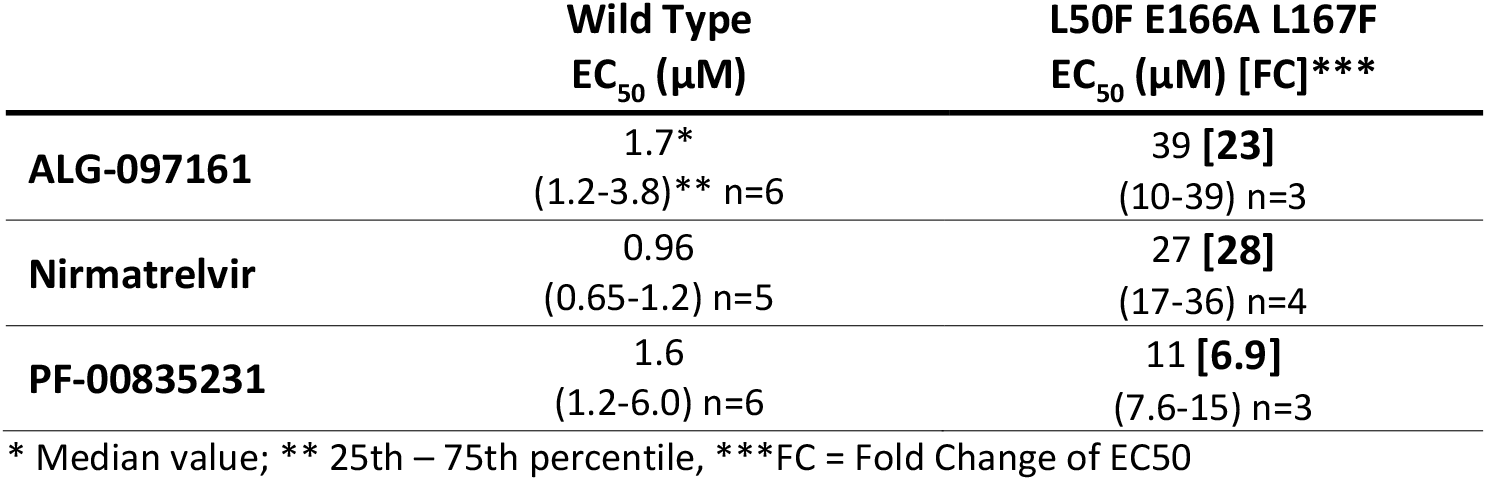
Phenotypic resistance associated with the L50F E166A L167F mutation profile as determined in a cell-based 3CLpro reporter assay.

|  | Wild Type<br>EC<sub>50</sub> (μM) | L50F E166A L167F<br>EC<sub>50</sub> (μM) [FC]*** |
| --- | --- | --- |
| **ALG-097161** | 1.7*<br>(1.2-3.8)** n=6 | 39 [23]<br>(10-39) n=3 |
| **Nirmatrelvir** | 0.96<br>(0.65-1.2) n=5 | 27 [28]<br>(17-36) n=4 |
| **PF-00835231** | 1.6<br>(1.2-6.0) n=6 | 11 [6.9]<br>(7.6-15) n=3 |

\* Median value; \*\* 25th – 75th percentile, \*\*\*FC = Fold Change of EC50

### Effect of amino acid substitutions on 3CLpro enzymatic activity

Recombinant 3CLpro proteins were produced with the L50F, E166A and L167F substitutions alone or combined, and their enzymatic activity was tested in a FRET assay using a peptide substrate featuring the canonical glutamine cleavage site [27]. In this assay, wild-type (WT) 3CLpro shows linear product conversion in the enzyme concentration range of 0.5-12.5 nM (Figure 3A). Compared with the WT protein, all tested mutants display reduced enzymatic activity (Figure 3B). The E166A and L167F, as single mutants, reduce the activity to about 12-20% of wild-type, whereas the L50F substitution reduces the activity to as low as 0.4%. The enzyme containing the three substitutions L50F E166A L167F is less compromised than L50F alone, with 4.6% activity as compared with WT. The loss in enzymatic activity for each mutant might be attributed to an apparent reduced binding affinity for the substrate, a loss in protein stability and/or a reduced formation of active dimers (Figure 4). The enzymes with E166A and L167F are less impaired than the enzyme with L50F. In comparison, the triple mutant exhibits an intermediate phenotype for both parameters (Figure 4).

**Figure 3:**
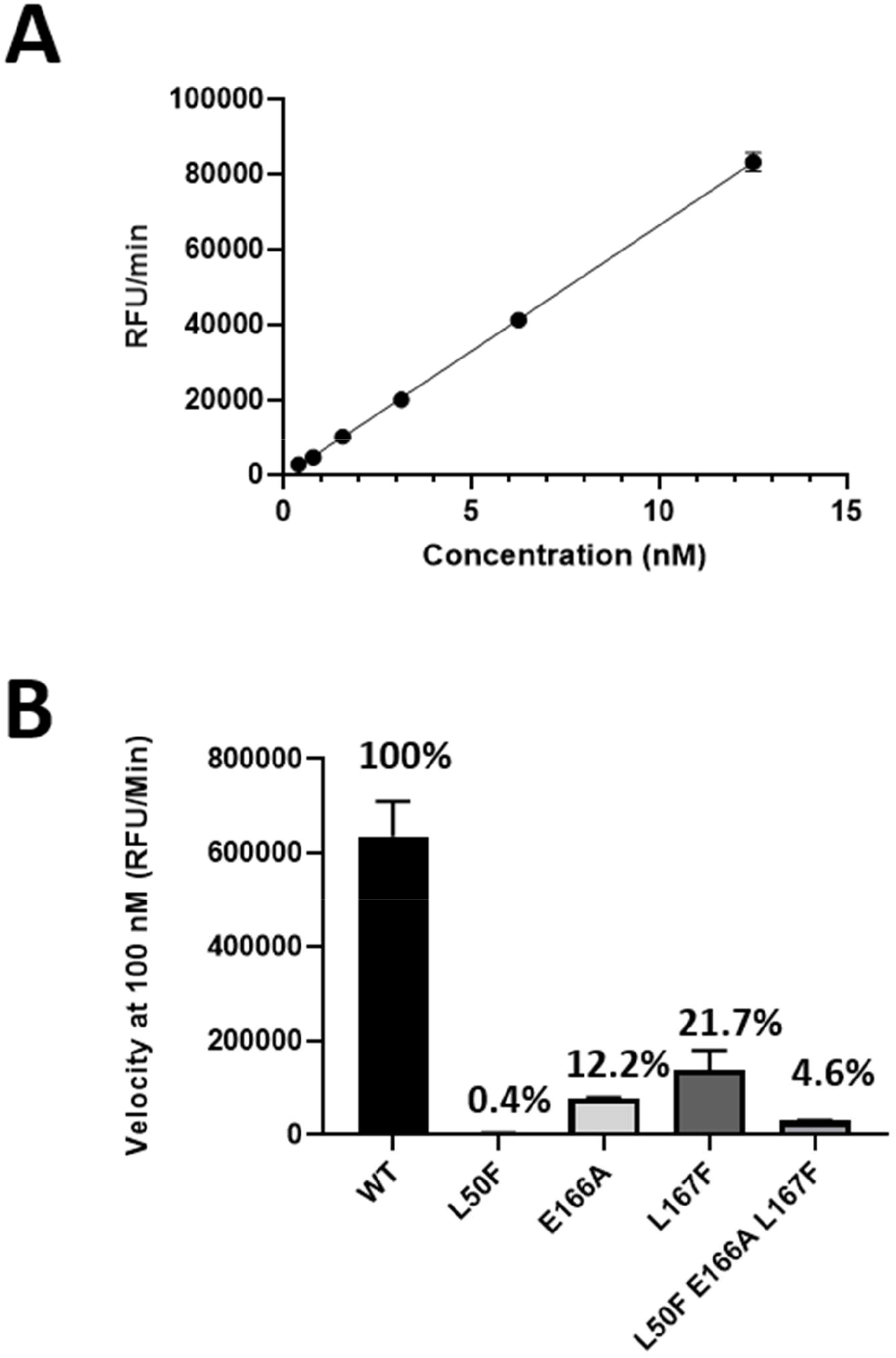
Enzymatic activity of WT and mutated SARS-CoV-2 3CLpro. (A) The enzymatic activity of WT 3CLpro was measured in a FRET assay. Three independent experiments were performed. The figure shows the results of one representative experiment. (B) Comparison of enzymatic activity between WT 3CLpro and mutated enzymes. Each data point represents the average of three independent experiments and mean and standard deviations are shown.

**Figure 4:**
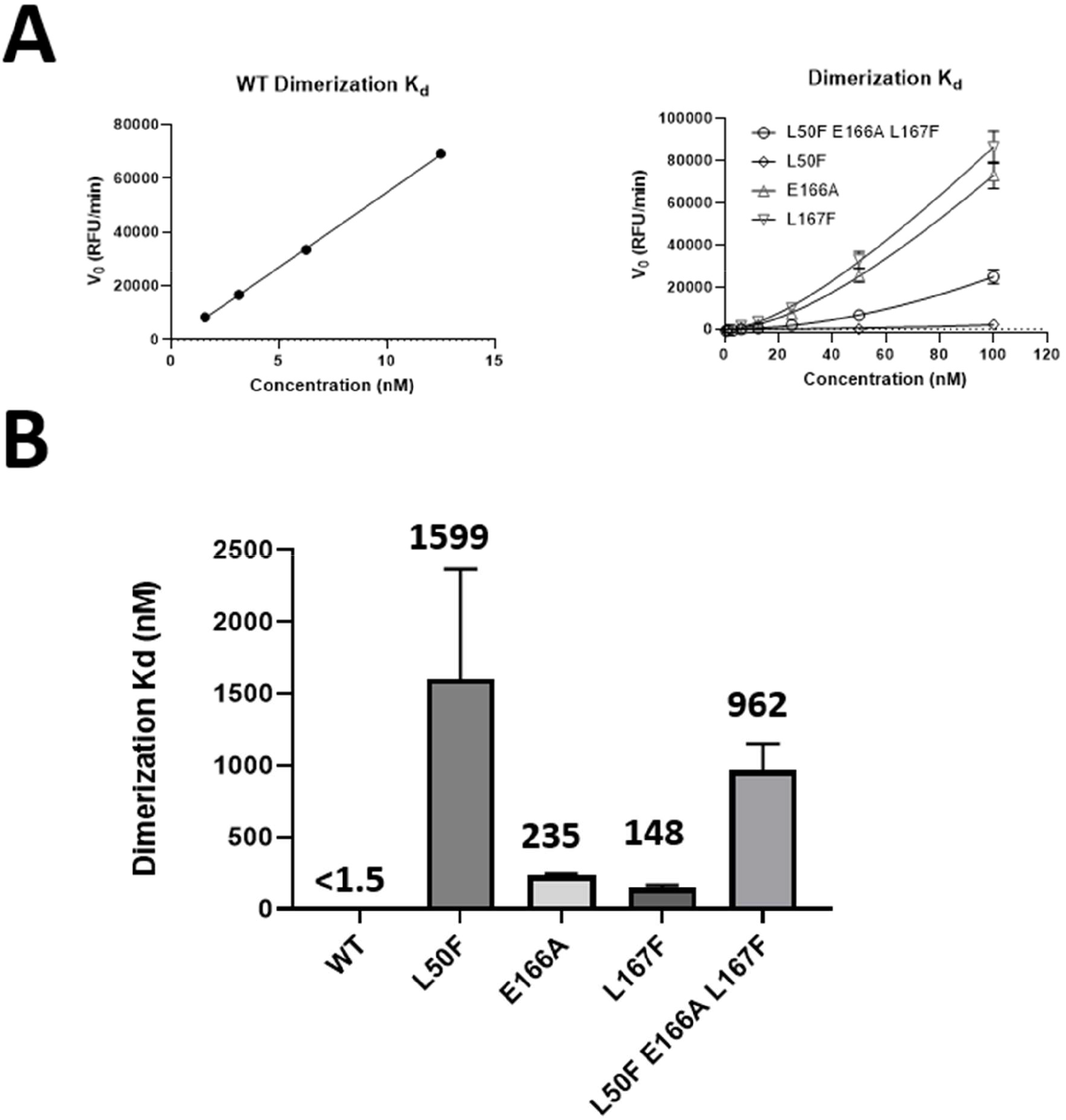
Effect of mutations on 3CLpro dimerization. (A) Initial velocities of enzyme titration of 3CLpro WT and mutant proteins were fitted as described (see methods) to calculate the monomer dimer equilibrium dissociation constant. Three independent experiments were performed. The figures show the results of one representative experiment. (B) 3CLpro dimerization binding affinity for each enzyme. Each data point represents the average of three independent experiments and mean and standard deviations are shown.

Ligand-induced protease activation and dimerization with low concentration of active site inhibitor has previously been described for MERS 3CLpro [28]. This is explained by the inhibitor stabilizing dimer formation by occupying only one of the two binding sites, while full occupancy of the two dimer active sites results in complete enzyme inhibition (Figure 5B). Although we did not perform direct protein dimerization studies, we could demonstrate that the impaired enzymatic activity of the mutants can be partially rescued with low concentrations (around 0.05 μM) of ALG-097161, PF-00835231 or nirmatrelvir, while higher concentrations inhibit product formation (Figure 5A; Supplemental S3).

**Figure 5:**
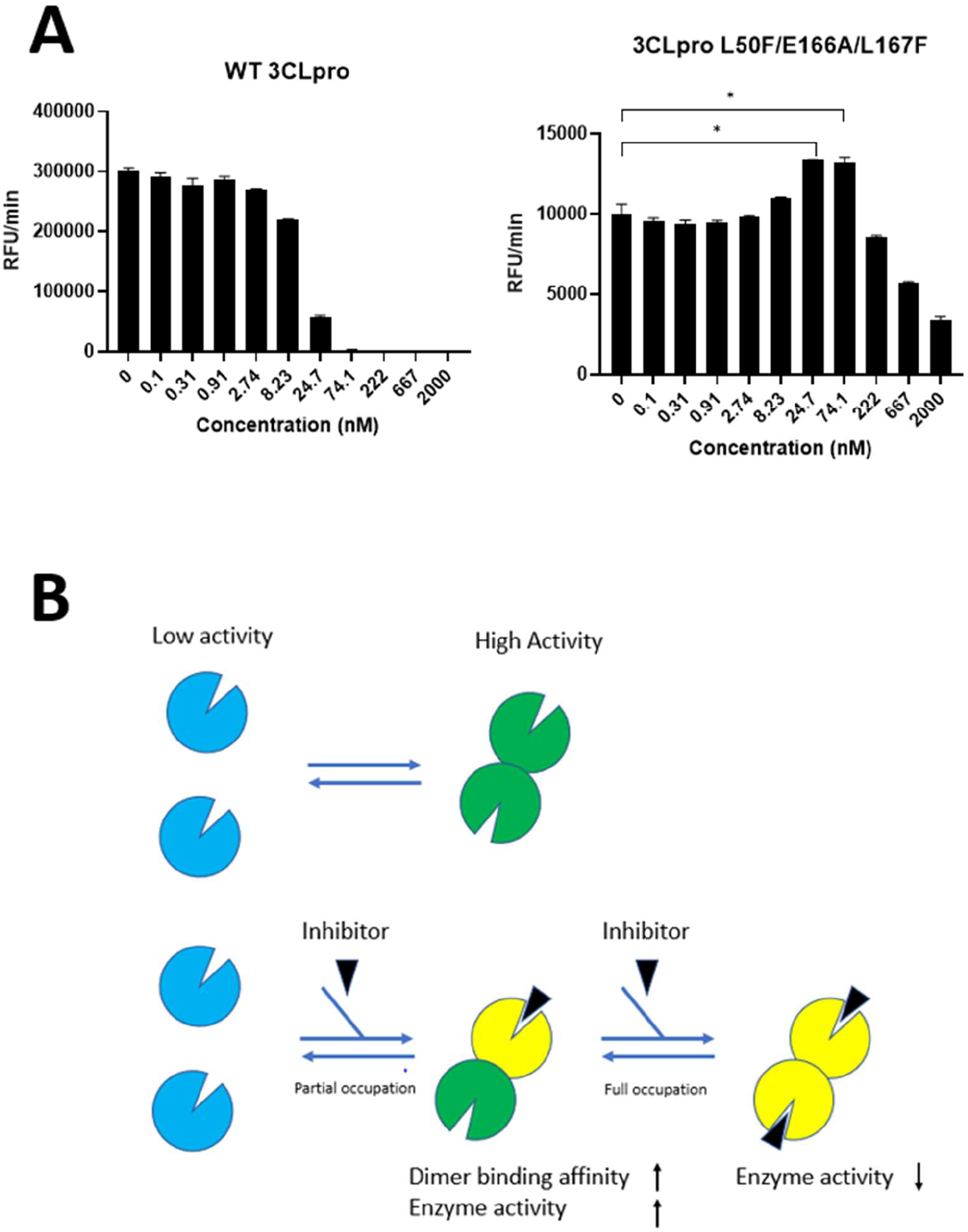
Ligand-induced enzymatic activation. (A) Comparison of ALG-097161 dose-response for WT and L50F E166A L167F 3CLpro. Three independent experiments were performed and mean and standard deviations are shown. The figure shows the results of one representative experiment. (* P< 0.05) Similar experiments with Nirmatrelvir and PF-00835231 are shown in Supplemental S3. (B) Model for ligand-induced dimerization and enzymatic activation at low concentration of inhibitor.

### Resistance to 3CLpro inhibitors in the biochemical assay

Based on the characterization of the different enzymes (see above), we decided to use all enzymes at a concentration of 50 nM to quantify the activity of the inhibitors. The effect of the L50F substitution could not be determined under these conditions due to its low intrinsic enzymatic activity. ALG-097161, nirmatrelvir and PF-00835231 all inhibit WT 3CLpro with IC_50_ values ranging from 13-23 nM (Table 4). For ALG-097161, the IC_50_ increases 5x-6x with single substitutions E166A and L167F, and reaches 35x for the enzyme with L50F E166A L167F. In comparison, the potency of nirmatrelvir is more affected by substitution E166A vs. L167F (10x vs. 4.4x) and reaches 72x increase on the enzyme with the 3 substitutions. For PF-00835231, the shift in IC_50_ is in general lower with a 6.0x change vs. the enzyme with L50F E166A L167F. However, the shift in IC_50_ is higher (∼100x) with the non-covalent and chemically distinct inhibitor ensitrelvir.

**Table 4:**
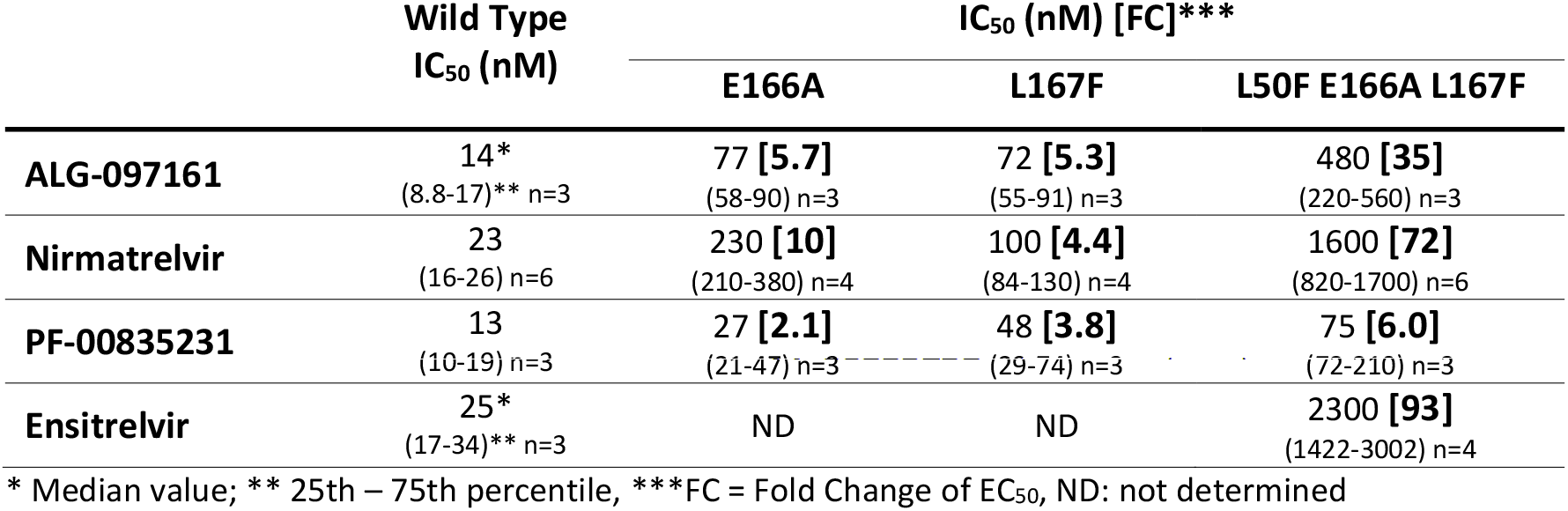
Biochemical analysis of the resistance associated with different substitutions in the 3CLpro enzyme.

### Structural Biology and Computational Chemistry

Experimental data with the 3CLpro system and computational chemistry investigations were used to rationalize the phenotype observed for ALG-097161 *in vitro*.

PF-00835231 and the associated crystal structure 6xhm provided a convenient starting point for modeling ALG-097161 in the catalytic pocket of 3CLpro, due to the structural similarity between both compounds. In particular, both structures feature a hydroxymethyl-ketone warhead that binds covalently to Cys145, as well as a 4-methoxyindole-P3 substituent.

Multiple conformations of ALG-097161 were used as input for docking calculations. These yielded a well-converged predicted binding mode of ALG-097161 covalently bound to the catalytic site of WT 3CLpro (Figure 6). In line with the crystal structures disclosed for similar compounds (e.g., 6xhm), ALG-097161 is predicted to form a covalent bond from the activated carbon atom of its hydroxymethyl-ketone warhead to the side chain of the catalytic C145 residue, anchoring the inhibitor into the binding cleft of 3CLpro. ALG-097161 forms seven direct hydrogen bonds with the SARS CoV2 3CLpro catalytic site. The oxygen atom of the inhibitor’s warhead interacts with the backbone NH of C145 in the oxyanion hole. The carbonyl oxygen of the P1 lactam hydrogen-bonds to the side chain of H163, while the NH of the P1 lactam is involved in a bifurcated interaction with the side chain of E166 and the backbone carbonyl of F140. The peptide backbone of ALG-097161 anchors itself with hydrogen bonds to the backbone carbonyl of H164 and the backbone HN of E166. The NH function of the P3 indole hydrogen-bonds the inhibitor to the backbone carbonyl of E166. The fused bicyclic P2 substituent of ALG-097161 maximizes Van der Waals interactions via extensive productive contacts to the protein side chains that form the S2 sub-pocket of WT 3CLpro. A key difference with PF-00835231 resides in the (lack of) interaction between the inhibitor backbone and Gln189; a direct consequence of the cyclization on P2, which results in the loss of a hydrogen bond donor site.

**Figure 6:**
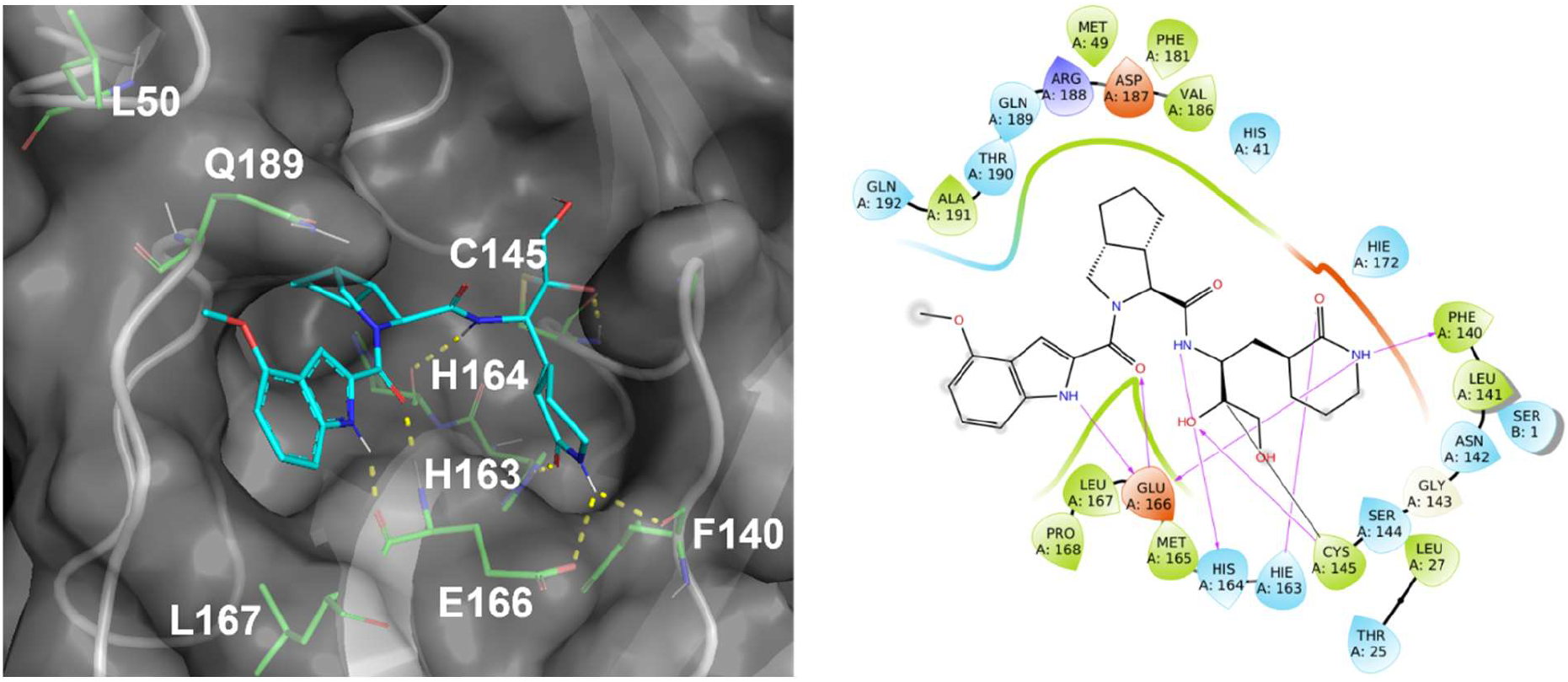
Predicted binding mode of ALG-097161 covalently bound to the catalytic site of WT 3CLpro. Left panel: ALG-097161 (carbon: cyan, nitrogen: navy; oxygen: red) covalently docked to WT 3CLpro (gray cartoon and surface; key side chains shown with carbon: green, nitrogen: navy; oxygen: red). Right panel: ALG-097161 binds covalently to the catalytic C145 and forms seven hydrogen bonds with 3CLpro: warhead to C145 in the oxyanion hole; P1 lactam to H163, E166 and F140; peptide backbone to H164 and E166; P3 indole to E166. The fused bicyclic P2 substitution maximizes Van der Waals interactions in the S2 sub-pocket.

The predicted binding mode of ALG-097161 within 3CLpro provides some rationale for the amino acid substitutions associated with resistance. In particular, residues 166 and 167 are located within 5Å of the bound ALG-097161. This observation can be extended to both PF-00835231 and nirmatrelvir, based on crystal structures from PDB codes 6xhm and 7rfw, respectively. Consequently, the observed substitutions have a direct impact on the compound /target interaction.

E166 plays a key role in compound binding, with no less than three hydrogen bonds formed with the inhibitors. The E166A substitution eliminates one of the hydrogen bonds made within the P1 sub-pocket. Interactions with this sub-pocket are important for substrate recognition by coronaviral 3CLpro enzymes, and its occupancy, in many cases by a lactam moiety, is a key driver for inhibitor potency. It is therefore unsurprising that all three inhibitors are adversely affected by the E166A change. Meanwhile, L167F results in the presence of a bulkier side chain on the neighboring position. While this change does not directly conflict with compound binding, the side chain of L167 is in close contact with other residues, including Phe185. Molecular Dynamics (MD) simulations on the L50F E166A L167F enzyme suggest that the bulkier Phe side chain results in some distortion of the neighboring sub-pocket. In particular, the loop containing L167F moves away from F185 and the distance between the Cαs of F185 and L167F / P168 consistently increases, resulting in a more open binding site and a sub-optimal fit of the inhibitor, even though H-bond interactions (including the two remaining H-bonds with the A166 backbone) appear conserved throughout 100ns simulations (Figure 7). Interestingly, all three inhibitors (ALG-097161, PF-00835231 or nirmatrelvir) were affected by the L167F change.

**Figure 7:**
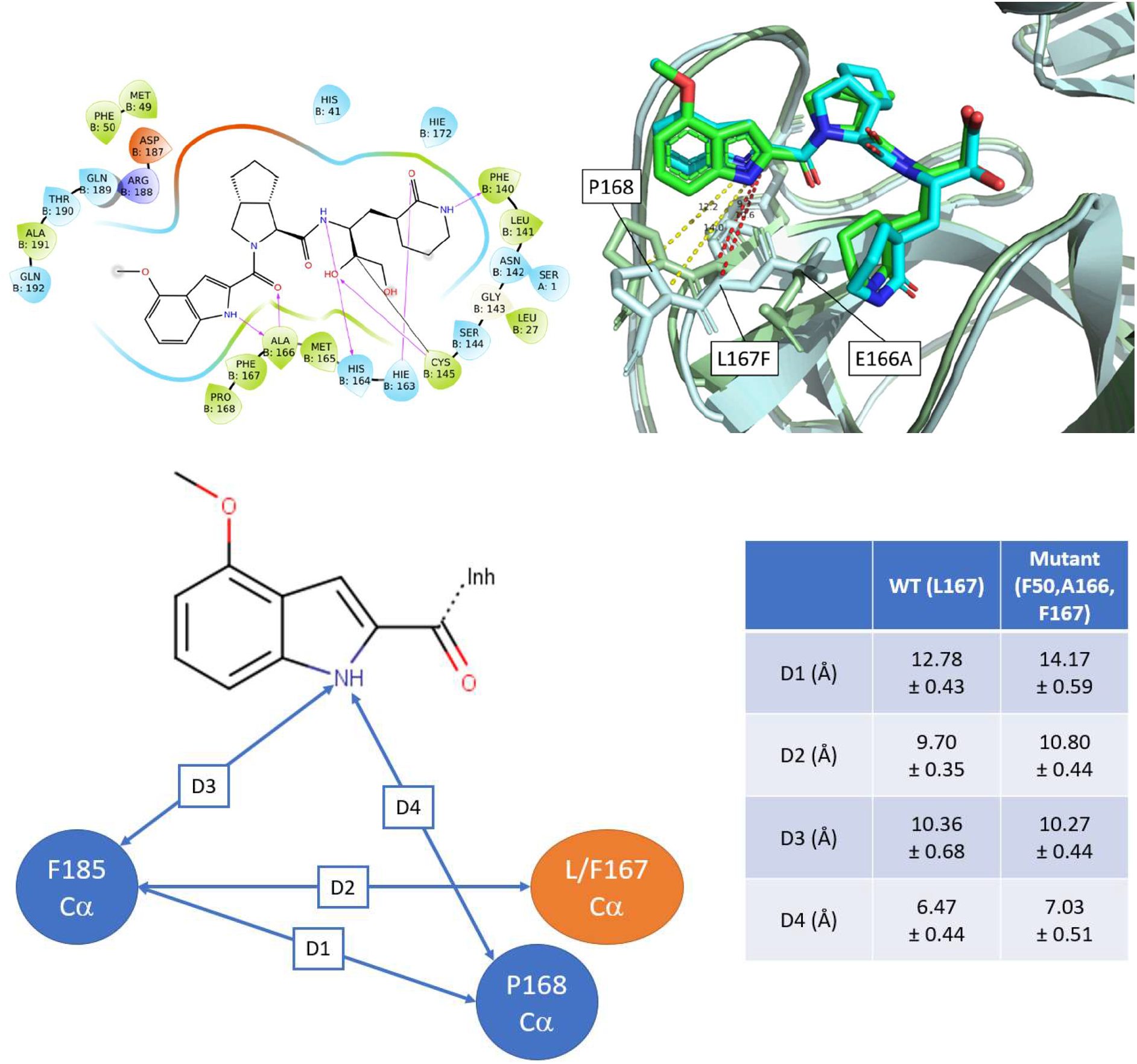
Proposed effect of the L167F substitution. Accommodation of the bulkier F167 residue results in some distortion of the binding site and in a possibly sub-optimal fit. Top Left: Interaction diagram for ALG-097161 in triple-mutant (L50F, E166A, L167F) 3CLpro. Interaction between the lactam moiety and the side-chain of residue 166 is lost due to mutation, while other direct H-bond interactions are conserved (>90% occurrence) over a 100 ns MD run. Top right: Overlay of representative MD frames for WT and mutant 3CLPro. WT protein displayed as pale green cartoon and L50F E166A L167F mutant displayed as pale blue cartoon. Bottom: Average distances between F185 (Cα), L167/F167 (Cα) P168 (Cα) and inhibitor. Distances collected over 2 x 380 frames (5ns → 100ns simulated time). Distances between F185 and L/F167 + P168 are increased following the L167F change.

Reasons behind the L50F mutation remain more elusive, as this residue falls outside of direct Van der Waals or hydrogen-bond contact range to ALG-097161. However, the L50 side chain is located only 2.7 Å from Q189, a residue which can interact with the peptide backbone of this inhibitor class either directly (as with PF-00835231) or *via* water molecules. L50F might therefore indirectly affect compound binding by altering the position of Q189 and the subsequent interactions mediated by this residue. However, a rigorous assessment of how this change impacts enzyme activity and/or inhibitor binding would require more elaborated techniques (e.g., free energy perturbation) that are beyond the purpose of this work.

Finally, it is anticipated that the natural SARS CoV-2 polyprotein substrates, that are processed by the 3CLpro, are less affected by the L50F, E166A and/or L167F substitutions, due to their larger size and approximately two-fold higher number of polar interactions with the protease (as observed in 2q6g.pdb for SARS CoV-1 3CLpro in complex with bound substrate).

## Discussion

Nirmatrelvir is the first oral SARS-CoV-2 (3CLpro) protease inhibitor granted emergency use authorization by the US FDA for the treatment of COVID-19. To avoid resistance development, HIV and HCV protease inhibitors are used, with great success, in fixed dose combinations with other directly acting antiviral drugs that have a different mechanism of action [29]. As a single agent, several of these inhibitors select (rapidly) for drug-resistant variants. SARS-CoV-2 infections are typically acute in nature; hence treatment can be of short duration, which may largely reduce the likelihood that resistant variants develop. Yet, also for acute infections, the emergence of drug-resistant variants is a serious concern, certainly in case that such variants can efficiently transmit resulting in a loss of therapeutic options for patients. For example, the first-generation influenza drugs, amantadanes, are no longer recommended, as resistance to amantadine and rimantadine is widespread [20].

To monitor clinical drug-resistance, it is crucial to identify the mutations involved. To that end, we used the 3CLpro inhibitor ALG-097161, a probe compound in the context of a drug discovery program, to select for resistant variants *in vitro* by passaging SARS-CoV-2 in presence of increasing drug concentrations. Similar experiments have been performed with SARS-CoV-2 neutralizing antibodies and polymerase inhibitors [22, 30].

The selection process indicated that there is a significant barrier to resistance for the 3CLpro inhibitor ALG-097161. The first changes were observed only between passage 5 and 8 (20-30 days). These were the double substitution L50F E166A in 3CLpro, which then evolved to the triple substitution L50F E166A L167F. Phenotyping this triple mutant virus revealed a 63x increase in the EC_50_ for ALG-097161. Moreover, significant cross-resistance (>10x increase EC_50_) was observed with nirmatrelvir, the active component of Paxlovid, and PF-00835231, an earlier 3CLpro clinical candidate. Next, we engineered an infectious clone with the only the 3CLpro substitutions L50F E166A L167F and the resistance against ALG-097161, nirmatrelvir and also ensitrelvir could be confirmed (>10x increase EC_50_). As a second confirmation, a cell-based assay was used with heterologous expression of wild-type and mutated 3CLpro (L50F E166A L167F) and a substrate linked to a reporter protein. The EC_50_ values for ALG-097161 and nirmatrelvir in this assay increase 23x and 28x respectively as compared with wild-type. Importantly, all of the viruses analyzed remained fully susceptible to the polymerase inhibitor remdesivir (GS-441524 form).

We further characterized the 3CLpro substitutions using biochemical assays. Enzymes carrying the single substitution L50F, E166A or L167F were found to have markedly lower enzymatic activity (<20% as compared to wild-type). In particular, the L50F mutant enzyme had only 0.4% activity compared to wild-type, which could be restored to 4.6% when also E166A and L167F were introduced. Each of the enzymes was also less efficiently inhibited by the 3CLpro inhibitors. The largest increase in IC_50_ was observed when all three substitutions L50F E166A L167F were combined. For this triple-mutant enzyme, the IC_50_ for ALG-097161, nirmatrelvir and ensitrelvir increases by 35x, 72x and 93x respectively.

A structural biology analysis reveals that the substitutions decrease interactions between the enzyme and the inhibitor. E166A causes a direct loss of a hydrogen bond with the lactam-moiety that is found in all the inhibitors tested here, L167F increases the size of the binding pocket causing a decrease of the Van der Waals forces between enzyme and inhibitor, and L50F is thought to change the rotamer conformation of Q189, which, for the wild-type, has many interactions with the inhibitors through a network of water molecules.

The loss of interactions, resulting from amino acid substitutions associated with resistance, might have a lower effect on substrates than on inhibitors, as the former rely on a significantly higher number of interactions for binding. Consequently, these resistance-associated mutations seem to allow a better discrimination between substrate and inhibitor at the expense of intrinsic enzymatic activity.

A recent sequence analysis across 4.9 million global SARS-CoV-2 isolates, circulating prior to the introduction of Paxlovid, showed a high genetic conservation of the 3CLpro protein [31]. From the 305 amino acid positions, only 3 have a polymorphism in >1% of the samples and 13 have a change in 1% - 0.11% of the samples. Changes at positions L50, E166 or L167 have a <0.11% prevalence. E166 displays extreme high conservation in this dataset with <0.001% samples having an alternative amino acid.

At this moment, peer-reviewed information on resistance against 3CLpro inhibitors in COVID-19 patients is, to our knowledge, not available. The Paxlovid label indicates that E166V is more common in nirmatrelvir/ritonavir-treated subjects relative to placebo-treated subjects (0.8% and <0.2% respectively), and in one patient, with a baseline L50F substitution, the E166V substitution co-occurred with L50F on Day 5 of treatment [32]. Both these observations are in accordance with the results described above.

While we can clearly observe resistance of different 3CLpro inhibitors for the L50F E166A L167V virus, we could not fully characterize the contribution of each substitution by itself. During the selection experiment, L50F together with E166A is selected first and L167V is acquired later. Additional cell-based experiments demonstrate that an L50F carrying virus is not resistant, while biochemical experiments indicate that E166A and L167F are associated with resistance. Taken together, we hypothesize that E166A is the main driver for resistance, that L50F is needed to support E166A and that presence of L167F further increases resistance. It would require an extensive study of the replication capacity of the different variants to support this hypothesis. Importantly, clear nirmatrelvir/ensitrelvir cross-resistance is observed with both the E166A L167F and L50F E166A L167F viruses.

During the writing of this manuscript, two preprints reported the selection of nirmatrelvir resistant SARS-CoV-2 viruses. One group performed > 480 independent selection experiments, resulting in the identification of multiple pathways of nirmatrelvir resistance [33]. They conclude that E166V confers the highest level of resistance of all substitutions tested (100x for nirmatrelvir; 23x for ensitrelvir). In addition, these authors show that E166V causes a reduced fitness that is restored by adding L50F or T21I. The selection of L50F E166V by nirmatrelvir *in vitro* has also been reported by another group [34]. Also, these authors come to the conclusion that E166V is responsible for resistance whereas L50F is a compensatory substitution required for maintaining fitness. As E166A and E166V are very similar substitutions (alanine and valine have similar chemical structures), these findings are in strong agreement with our observations. A third group engineered a chimeric VSV-variant, that has SARS-CoV-2 3CLpro expressed in an artificial polyprotein, for which the replication depends on 3CLpro activity. Selection for nirmatrelvir resistance resulted in the appearance of L167F in the absence of other mutations, and an engineered SARS-CoV-2 infectious clone with only the L167F substitution showed a marginal level of resistance for nirmatrelvir (2x increase EC_50_)[35].

Important to mention is also the work of Flynn et al [36], who performed different functional 3CLpro screens in yeast to study the resilience of all amino acid positions towards substitutions. Interestingly, the authors conclude that position E166 is highly tolerant to substitutions including the E166A or E166V substitution. Based on available structural biology data of 3CLpro in complex with inhibitors or natural substrate and the functional screen data, these authors identified E166A and E166Q as potentially resistance associated substitutions.

Our report emphasizes the need for additional research to elucidate potential resistance pathways of SARS-CoV-2 inhibitors *in vitro*. It is also a starting point for the surveillance of nirmatrelvir resistance and indicates the potential need for combination therapies with different classes of SARS-CoV-2 inhibitors.

## Methods

### Compounds, cells, viruses, proteins and peptides

ALG-097161, Nirmatrelvir and PF-00835231 were synthesized by Aligos Therapeutics and purified to >95% purity. GS-441524 was obtained from MedChem Express (cat no HY-103586). The African green monkey kidney Vero E6 cell line was purchased from ATCC (catalog no. CRL-1586™) and maintained in DMEM (gibco cat no 41965-039) supplemented with 10% v/v heat-inactivated FCS. The SARS-CoV-2 GHB-03021 (EPI ISL407976|2020-02-03) isolate was obtained from a Belgian patient returning from Wuhan in February 2020. The isolate was passaged 7 times on VeroE6 cells, which introduced two series of amino acid deletions in the spike protein [37].

SARS-CoV-2 3CLpro wild-type and mutant enzymes were produced as previously described [38]. Peptide substrate (Dabcyl-KTSAVLQSGFRKM-E(Edans)-NH_2_ for FRET was sourced from Biopeptide (San Diego, CA) at >95% purity.

### Genotypic analysis

#### Illumina sequencing

RNA was extracted from cell culture supernatant, using the NucleoSpin RNA virus kit (Macherey-Nagel) according to the manufacturer’s instructions. Whole genome sequencing was outsourced to EurofinsGenomics (ARTIC SARS-CoV-2 WGS, Konstanz, Germany), who performed reverse transcription, enrichment of the viral genome using a primer set similar to the ARTIC primers (> 200 primer pairs, covering the full 29.9 kb viral genome), generation of libraries, Illumina sequencing (2x150 bp read mode), and sequence cleaning to remove adapters and poor-quality bases. Sequences were further analyzed using Geneious Prime software (v2022.2.1) by mapping to the SARS-CoV-2 RefSeq NC_045512, and variant calling as described by the software manufacturer (https://help.geneious.com/hc/en-us/articles/360045070991-Assembly-of-SARS-CoV-2-genomes-from-tiled-amplicon-Illumina-sequencing-using-Geneious-Prime). For each sample, >0.9 M reads that could be aligned with the SARS-CoV-2 genome were obtained with >99.5% of untrimmed bases being of high-quality, and a mean coverage of > 4000 reads.

#### Oxford Nanopore sequencing

As a control, the same samples were also sequenced using an inhouse pipeline. For this Reverse transcription was carried out via SuperScript IV and cDNA was posteriorly amplified using Q5® High-Fidelity DNA Polymerase (NEB) with the ARTIC nCov-2019 primers, following the recommendations in the sequencing protocol of the ARTIC Network (https://artic.network/ncov-2019). Samples were multiplexed following the manufacturer’s recommendations, using the Oxford Nanopore Native Barcoding Expansion kits NBD104 (1-12) and NBD114 (13-24), in conjunction with Ligation Sequencing Kit 109 (Oxford Nanopore). Sequencing was carried out on a MinION sequencer using R9.4.1 flow cells and MinKNOW 2.0 software. Both methods resulted in the same consensus sequence for each sample. For further mutation analysis we used the deep sequencing data generated by the Illumina protocol.

### Antiviral testing

The 50% effective concentration (EC_50_), the concentration of compound required for fifty percent antiviral activity, was determined on VeroE6 cells as follows. On day -1, the test compounds were serially diluted in 100 μL assay medium (DMEM supplemented with 2% v/v FCS) in 96-well plates. In a next step, 50 μL of VeroE6 cells (25,000 cells/well) was added together with 2 μM of the MDR1-inhibitor CP-100356 (final concentration 0.5 μM). The plates were incubated (37°C, 5% CO2 and 95% relative humidity) overnight. On day 0, 50 μL of SARS-CoV-2 WT or the triple mutant strain at a multiplicity of infection (MOI) of 0.001 tissue culture infectious dose (TCID50) per cell was added and the plates were stored in a humidified incubator at 37°C and 5% CO2. In absence of antiviral activity, the VeroE6 cells undergo a cytopathic effect. Cell viability was determined 4 days p.i. using a viability staining with MTS [39]. The percentage of antiviral activity was calculated by subtracting background and normalizing to the untreated-uninfected control wells, and the EC_50_ was determined using logarithmic interpolation.

### Generation and rescue of recombinant viruses

We used the in-yeast transformation-associated recombination (TAR) cloning method as described previously with some adaptations [25]. In sum, the whole SARS-CoV-2 genome was encoded in 12 overlapping DNA fragments. These so-called WU-Fragments and a TAR-vector are homologous recombined in yeast forming the yeast artificial chromosome (YAC). The SARS-CoV-2 3CLpro is encoded on WU-Fragment 5 and was replaced with newly generated and overlapping PCR products to introduce the amino acid changes L50F, E166A L167F or L50F E166A L167F. The overlapping PCR products were made via reverse transcription from RNA purified from vRNA of the virus stock obtained from the selection. In brief, cDNA was generated from RNA by LunaScript RT SuperMix (NEB). PCR reactions targeting the 3CLpro gene were performed using Q5® High-Fidelity DNA Polymerase (NEB) and the resulting PCR products were mixed and matched to recombine SARSCoV-2 3CLpro^L50F^, SARSCoV-2 3CLpro^E166A L167F^ and SARSCoV-2 3CLpro^L50F E166A L167F^. All PCR products were purified with the High Pure PCR Product Purification Kit (Roche) before being used for TAR cloning.

In vitro transcription was performed for EagI-cleaved YACs and PCR-amplified SARS-CoV-2 N gene using the T7 RiboMAX Large Scale RNA production system (Promega) as described previously [25]. Transcribed capped mRNA was electroporated into baby hamster kidney (BHK-21) cells expressing SARS-CoV N protein. Electroporated cells were co-cultured with susceptible Vero TMPRSS2 cells to produce passage 0 (P.0) of the recombinant viruses. Passage 1 virus stocks were further produced in Vero TMPRSS2 cells and the presence of mutations was confirmed by Illumina sequencing. These results also showed the absence of other non-synonymous mutations in the viral genome.

### 3CLpro FRET-based assay

The FRET-based assay was performed similarly to the SAMDI-MS assay. Assays were performed in 20 μL volume in 384-well Non-binding Low-volume plates (Greiner Bio-One; Monroe, NC) at ambient temperature. 3CLpro and its mutants were preincubated with inhibitors for 30 min. Reactions were initiated by the addition of a FRET-compatible peptide substrate Dabcyl-KTSAVLQSGFRKM-E(Edans)-NH_2_. Fluorescence was measured for 90 minutes at 2-min intervals using a 340/460 excitation/emission filters on a Envision plate reader (Perkin Elmer). The IC_50_ values were calculated by fitting the curves using a four-parameter equation in GraphPad Prism. To calculate the dimer dissociation constant (K_d_), the velocities of enzyme titration of WT and mutant 3CLpros were fitted to the equations (1) and (2).

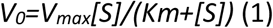

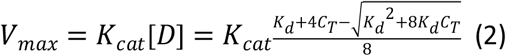

Equation (2) was described previously for the calculation of monomer dimer equilibrium dissociation constant (K_d_) [28].

### 3CLpro cell-based reporter assay

The cell-based reporter assay of SARS-CoV-2 3CLpro enzymatic function was described previously [26]. This gain-of-signal assay is based on the expression of a chimeric protein consisting of Src-Mpro-Tat-eGFP in which the Mpro amino acid sequence is flanked with cognate N- and C-terminal self-cleavage sites. While it was expected that Mpro inhibition would result in a different subcellular localization of the GFP signal, it was found that without Mpro inhibition there is no detectable GFP expression, while in the presence of Mpro inhibition there is high GFP expression. Hence the naming gain-of-signal assay. The exact mechanism behind this reporter system is not known but results suggest a mechanism in which Mpro activity somehow suppresses the accumulation of reporter mRNA in transfected cells.

293T cells were maintained at 37°C/5% CO2 in Dulbecco’s DMEM (Corning#10-013-CV) supplemented with 10% fetal bovine serum (Gibco #10091148) and 1% penicillin/streptomycin (Gibco #15140122). 293T cells were seeded in a 96-well plate at 5×10^5^ cells/well and returned to incubation for 24h. The following day, cells were treated with compound at a final DMSO concentration of 2% prior to transfection with 100 ng of the wild-type or triple mutant (L50F E166A L167F) plasmid with Lipofectamine LTX (ThermoFisher#15338100) for 24h. Post 24h transfection, GFP fluorescence was detected with a Victor (Perkin Elmer) using the Fluorescein 485/535 setting. Inhibition was determined from the resulting fluorescent signal and graphed using a variable slope four parameter curve fitting with GraphPad Prism (v 9.2.0).

### Modeling of inhibitors in WT and mutant (L50F E166A L167F) 3CLpro

PDB structure 6xhm (complex with PF-00835231) was used as starting point for modeling compounds in WT and mutant (L50F E166A L167F) 3CLpro. The receptor structure was cleaned up and minimized with restraints ahead of docking using the Protein Preparation Wizard in maestro (Schrödinger). Ligands were processed using the Ligprep module in maestro. Stereoisomers were not generated, as the absolute configuration of the ligand was known.

Covalent docking was used to predict the binding mode of ALG-097161. C145 was specified as the reactive residue for covalent binding, and the centroid of the X-ray reference ligand was used to define the investigated binding site. A custom covalent reaction parameter file was used to define the reaction between the warhead moiety and the catalytic C145. High accuracy (Pose Prediction Thorough) docking methodology was implemented for pose prediction. Initial docking poses within 4 kcal/mole (glide gscore) were collected for further processing. Residues within 6 Å of the bound compound were minimized after covalent bond formation. The top 5 binding mode suggestions were saved for visual inspection.

Initial structures for the L50F E166A L167F triple mutant were built by introducing the mutations directly into the WT complex. An initial restrained minimization was done with the protein preparation wizard to remove any unacceptable clash. Next, a Desmond simulation system was built (see below) and a 100 ps energy minimization procedure was run with Desmond to reach the nearest energy minimum.

Molecular Dynamics (MD) simulations were performed using the Desmond-GPU package. A simulation system was first set up around the protein-inhibitor complex, including a 10 Å water box, Na+ ions necessary to neutralize the system and additional Na+ and Cl-ions to simulate a 0.15 M NaCl concentration. OPLS4 was selected as force field. The NPT ensemble (T = 300K, P = 1.01325 bar) was used. The system was relaxed prior to simulation. MD were run for a (simulated) period of 100 ns, with frames saved every 0.25 ns (total 400 frames). Final frames were energy-minimized (100 ps minimization protocol). Protein-ligand interactions and distances were evaluated in the 5 ns - 100 ns period.

## Acknowledgments

We thank Tina Van Buyten and Niels Cremers for their excellent technical assistance and Myriam Cornelis for dedicated editorial assistance.

