## Supplemental for "The substitutions L50F, E166A and L167F in SARS-CoV-2 3CLpro are selected by a protease inhibitor *in vitro* and confer resistance to nirmatrelvir"

**Supplemental S1: Enzymatic data against 3CLpro and Cathepsin L**

| | $IC_{50}$ (nM) | |
|---|---|---|
| | **3CLpro** | **Cathepsin L** |
| **ALG-097161** | 14*<br>(8.8-17)** n=3 | > 10,000<br>n=1 |
| **Nirmatrelvir** | 23<br>(16-26) n=6 | > 10,000<br>n=1 |
| **PF-00835231** | 13<br>(10-19) n=3 | 130<br>(61 – 260) n=58 |
| **Ensitrelvir** | 25*<br>(17-34)** n=3 | > 10,000<br>n=1 |

* Median value
** 25th – 75th percentile
***Cathepsin L assay was performed as described previously (Liu, C., et al., Antiviral Res, 2021. 187: p. 105020.). In this endpoint assay the signal was measured at 30 min after initiation.

**Supplemental S2: Genotypic analysis**

Viral RNA was extracted from the supernatant of virus cultures, containing ALG-097161, at passage 5, 8 and 12. Samples were analyzed by ARTIC SARS-CoV-2 whole genome sequencing (EurofinsGenomics, Germany). For each sample >0.9 M reads were obtained that could be aligned with the SARS-CoV-2 genome.
The sequence information of each virus sample was compared with the consensus sequence of the starting SARS-CoV-2 GHB-03021 strain. Information of all nonsynonymous nucleotide changes, that were present in > 10% of the reads of any of the samples, are shown in the table below.

| Nucleotide Number (NC_045512) | Change | Gene | CDS Codon Number | Gene Product | 3CLpro Codon Number | Codon Change | Amino Acid Change | Variant Frequency | | |
|---|---|---|---|---|---|---|---|---|---|---|
| | | | | | | | | Passage 5 | Passage 8 | Passage 12 |
| 1547 | A -> C | ORF1ab | 428 | nsp2 | - | AGC -> CGC | S -> R | < 10% | 10.4% | 10.0% |
| 7252 | G -> U | ORF1ab | 2329 | nsp3 | - | UUG -> UUU | L -> F | 17.7% | 18.0% | 18.1% |
| 7254 | C -> G | ORF1ab | 2330 | nsp3 | - | GCA -> GGA | A -> G | 11.5% | 11.6% | 11.8% |
| 10202 | C -> U | ORF1ab | 3313 | nsp5_3CLpro | 50 | CUU -> UUU | L -> F | < 10% | 99.5% | 99.8% |
| 10551 | A -> C | ORF1ab | 3429 | nsp5_3CLpro | 166 | GAA -> GCA | E -> A | < 10% | 88.1% | 99.6% |
| 10555 | A -> U | ORF1ab | 3430 | nsp5_3CLpro | 167 | UUA -> UUU | L -> F | < 10% | < 10% | 98.2% |
| 15671 | A -> G | ORF1ab | 5136 | nsp12 | - | GAG -> GGG | E -> G | 58.1% | < 10% | < 10% |
| 16370 | U -> C | ORF1ab | 5369 | nsp13 | - | GUU -> GCU | V -> A | 34.5% | 97.7% | 99.0% |
| 16985 | C -> U | ORF1ab | 5574 | nsp13 | - | ACU -> AUU | T -> I | < 10% | 99.0% | 98.4% |
| 17057 | U -> A | ORF1ab | 5598 | nsp13 | - | AUG -> AAG | M -> K | 59.5% | < 10% | < 10% |
| 17332 | A -> G | ORF1ab | 5690 | nsp13 | - | ACG -> GCG | T -> A | < 10% | 80.3% | 98.6% |
| 21005 | C -> U | ORF1ab | 6914 | nsp16 | - | GCA -> GUA | A -> V | < 10% | < 10% | 99.2% |
| 21866 | A -> G | S | 102 | S | - | AGA -> GGA | R -> G | 37.9% | 99.7% | 99.7% |
| 24968 | A -> G | S | 1136 | S | - | ACA -> GCA | T -> A | < 10% | 38.8% | 97.6% |
| 25538 | G -> U | ORF3a | 49 | ORF3a | - | GGC -> GUC | G -> V | < 10% | 78.2% | 97.8% |
| 26267 | A -> U | E | 8 | E | - | GAG -> GUG | E -> V | 57.7% | < 10% | < 10% |

| Color Coding: | < 25% | 25% - 75% | 75% - 90% | > 90% |
|---|---|---|---|---|

Note that the sequences encoding the 3CLpro cleavage sites are at nucleotide position:
Nsp4-5: 10040-10066; Nsp5-6: 10958-10984; Nsp6-7: 11828-11854; Nsp7-8: 12077-12103; Nsp8-9: 12671-12697; Nsp9-10: 13010-13036; Nsp1011: 13427-13453; Nsp12-13:16222-16248; Nsp13-14: 18025-18051; Nsp14-15:19609-19632; Nsp15-16: 20644-20670

**Supplemental S3: Ligand-induced enzymatic activation.**

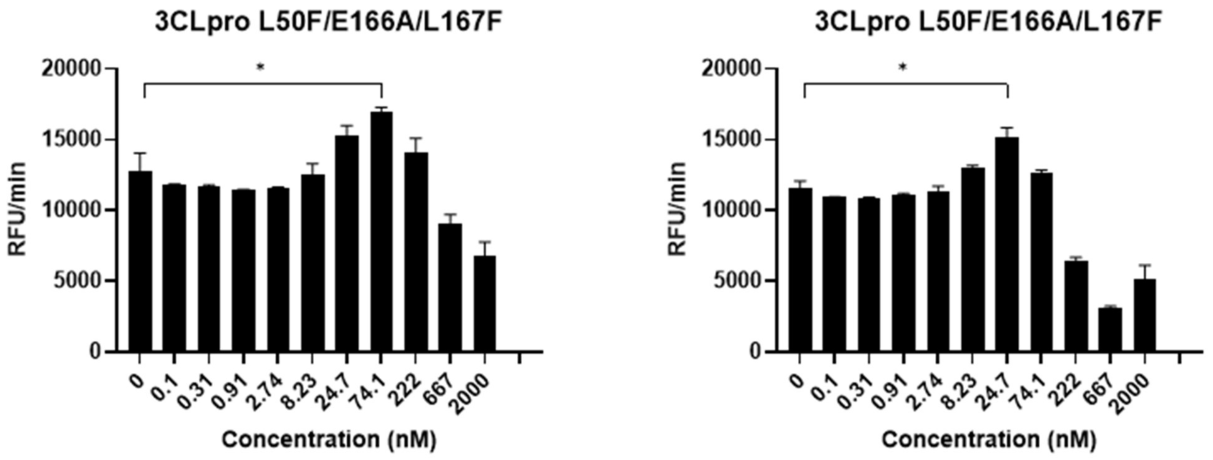


Nirmatrelvir PF-00835231

Figure S3: **Ligand-induced enzymatic activation.** Nirmatrelvir and PF-00835231 dose-response for L50F E166A L167F 3CLpro. (* P< 0.05). Supplemental to fig 5a of the manuscript.

### Supplemental S4: Chemical synthesis of ALG-097161

**Scheme S1. Synthesis of Intermediate 4[a]**

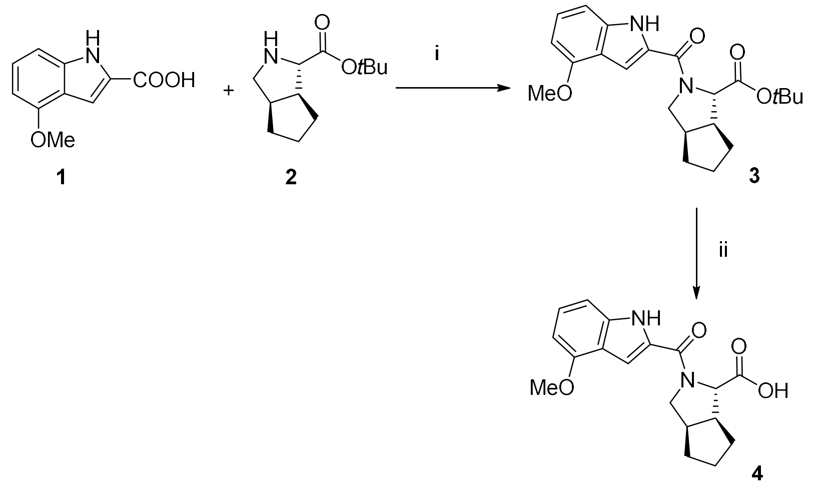


[a]Reagents and conditions: (i) HATU, DIPEA, DMF, 78%. (ii) TFA, $CH_2Cl_2$, quantitative.

***tert*-Butyl (1*S*,3a*R*,6a*S*)-2-(4-methoxy-1*H*-indole-2-carbonyl)octahydrocyclopenta[*c*]pyrrole-1-carboxylate (3)**

A mixture of 4-methoxy-1*H*-indole-2-carboxylic acid (348 mg, 1.82 mmol), HATU (693 mg, 1.82 mmol) and DIPEA (0.866 mL, 4.97 mmol) in DMF (5 mL) was stirred at rt for 30 min. *tert*-Butyl (1*S*,3a*R*,6a*S*)-octahydrocyclopenta[*c*]pyrrole-1-carboxylate oxalate (500 mg, 1.65 mmol) was added and the reaction mixture was stirred at rt for 3 h. The reaction mixture was diluted with water (20 mL) and was extracted with EtOAc (3 x 50 mL). The organic phases were combined, washed with brine (2 x 30 mL), dried over $Na_2SO_4$, filtered and concentrated under reduced pressure. The residue was purified by flash chromatography on silica gel using a gradient of EtOAc (0-30%) in petroleum ether to afford 530 mg (78%) of the title compound as a white solid. LCMS (ESI, m/z): 329 $[M-56+ H]^+$.

$^1H$ NMR (300 MHz, DMSO-$d_6$) δ 11.54 - 11.65 (m, 1H), 7.08 - 7.18 (m, 1H), 7.00 - 7.07 (m, 1H), 6.88 - 6.99 (m, 1H), 6.46 - 6.56 (m, 1H), 4.21 - 4.34 (m, 1H), 4.01 - 4.18 (m, 1H), 3.84 - 3.96 (m, 3H), 3.69 - 3.82 (m, 1H), 2.52 - 2.94 (m, 2H), 1.49 - 2.05 (m, 6H), 1.28 - 1.47 (m, 9H).

**(1*S*,3a*R*,6a*S*)-2-(4-methoxy-1*H*-indole-2-carbonyl)octahydrocyclopenta[*c*]pyrrole-1-carboxylic acid (4)**

To a solution of **3** (200 mg, 0.520 mmol) in $CH_2Cl_2$ (3 mL) cooled at 0°C was added TFA (1 mL). The reaction mixture was stirred at rt for 1 h and was concentrated under reduced pressure to afford quantitatively the title compound which was used in the next step without further purification. LCMS (ESI, m/z): 329 $[M+H]^+$.

**Scheme S2. Synthesis of Intermediate 9[a]**

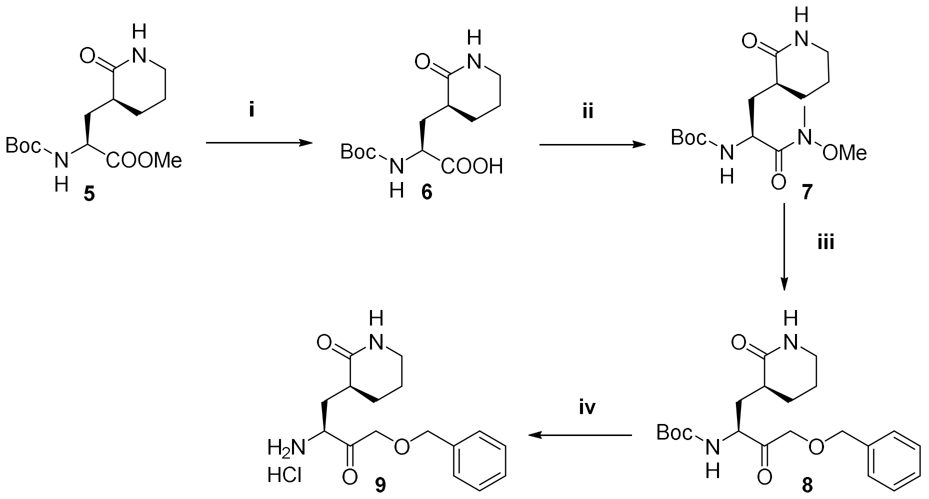


[a]Reagents and conditions: (i) 3M NaOH, MeOH, 60%. (ii) *N*,*O*-Dimethylhydroxylamine hydrochloride, NMM, HOBt, ECDI, $CH_2Cl_2$, 66%. (iii) Mg, $HgCl_2$, benzylchloromethyl ether, THF, 28%. (iv) 4N HCl in dioxane, dioxane, quantitative.

**(*S*)-2-((*tert*-Butoxycarbonyl)amino)-3-((*S*)-2-oxopiperidin-3-yl)propanoic acid (6)**

To a solution of methyl (*S*)-2-((*tert*-butoxycarbonyl)amino)-3-((*S*)-2-oxopiperidin-3-yl)propanoate (3.00 g, 10.0 mmol, synthesized according to *J. Med. Chem.*, **2015**, 9414-9420) in MeOH (15 mL) cooled at 0°C was added 3M NaOH (15 mL, 45.0 mmol). The reaction mixture was stirred at 0°C for 1 h. The reaction mixture was partially concentrated under reduced pressure to remove MeOH. The residue was extracted with EtOAc (3 x 30 mL). The organic phases were combined, washed with brine (2 x 20 mL), dried over $Na_2SO_4$, filtered and concentrated under reduced pressure to provide 1.70 g (60%) of the title compound as a white solid. . $^1$H NMR (300 MHz, DMSO-$d_6$) δ 12.49 (br s, 1H), 7.47 (s, 1H), 7.22 (d, *J* = 8.3 Hz, 1H), 3.88 - 4.05 (m, 1H), 3.09 - 3.11 (m, 2H), 2.07 - 2.16 (m, 2H), 1.82 - 1.96 (m, 1H), 1.68 - 1.83 (m, 1H), 1.57 - 1.62 (m, 2H), 1.38 - 1.40 (m, 10H). LC-MS (ESI, m/z): 287 $[M+H]^+$.

***tert*-Butyl ((*S*)-1-(methoxy(methyl)amino)-1-oxo-3-((*S*)-2-oxopiperidin-3-yl)propan-2-yl)carbamate (7)**

To a mixture of **6** (1.70 g, 5.93 mmol) in $CH_2Cl_2$ (25 mL) cooled at 0°C were added *N*,*O*-dimethylhydroxylamine hydrochloride (0.575 g, 5.93 mmol), NMM (1.80 g, 17.8 mmol), HOBt (0.800 g, 5.93 mmol) and EDCI (1.25 g, 6.53 mmol). The reaction mixture was stirred for 1 h at 0°C. Water (20 mL) was added and the phases were separated. The organic phase was washed with 1M HCl (2 x 20 mL), water (20 mL), sat. $NaHCO_3$ (2 x 20 mL) and brine (20 mL), dried over $Na_2SO_4$, filtered and concentrated under reduced pressure to provide 1.29 g (66%) of the title compound as a yellow solid. LC-MS (ESI, m/z): 330 $[M+H]^+$. $^1H$ NMR (300 MHz, DMSO-$d_6$) δ 7.42 (s, 1H), 7.13 - 7.15 (d, $J$ = 8.4 Hz, 1H), 4.44 - 4.50 (m, 1H), 3.72 (s, 3H), 3.05-3.20 (m, 5H), 1.99 - 2.24 (m, 2H), 1.85 - 1.97 (m, 1H), 1.69 - 1.83 (m, 1H), 1.58 - 1.69 (m, 1H), 1.37 - 1.44 (m, 11H).

***tert*-Butyl ((*S*)-4-(benzyloxy)-3-oxo-1-((*S*)-2-oxopiperidin-3-yl)butan-2-yl)carbamate (8)**

To a mixture of Mg (1.39 g, 57.3 mmol) and $HgCl_2$ (1.04 g, 3.83 mmol) in THF (50 mL) cooled at -45°C was added benzylchloromethyl ether (8.99 g, 57.4 mmol). The mixture was stirred at -45-5°C for 5 h. After cooling to - 45°C, **7** (2.10 g, 6.37 mmol) was added. The reaction mixture was slowly allowed to warm to rt and was stirred at rt overnight. The reaction was quenched by addition of sat. $NH_4Cl$ (30 mL). The reaction mixture was extracted with EtOAc (3 x 50 mL). The organic phases were combined, washed with brine (2 x 50 mL), dried over $Na_2SO_4$, filtered and concentrated under reduced pressure. The residue was purified by flash chromatography on silica gel using a gradient of MeOH (0-5%) in $CH_2Cl_2$ to afford 700 mg (28%) of the title compound as a yellow oil. LCMS (ESI, m/z): 391 $[M+H]^+$. $^1H$ NMR (400 MHz, Chloroform-$d$) δ 7.45 - 7.29 (m, 5H), 6.03 (s, 1H), 5.90 - 5.92 (m, 1H), 4.47 - 4.72 (m, 3H), 4.21 - 4.42 (m, 2H), 3.27 - 3.30 (m, 2H), 2.30 - 2.42 (m, 1H), 2.04 - 2.21 (m, 2H), 1.80 - 1.92 (m, 2H), 1.65 - 1.77 (m, 1H), 1.48 - 1.57 (m, 1H), 1.44 (s, 9H).

**(*S*)-3-((*S*)-2-Amino-4-(benzyloxy)-3-oxobutyl)piperidin-2-one hydrochloride (9)**

To a solution of **8** (300 mg, 0.768 mmol) in dioxane (3 mL) cooled at 0°C was added 4N HCl in dioxane chloride (3.0 mL, 12.0 mmol). The reaction mixture was stirred at rt for 1 h and was concentrated under reduced pressure to afford quantitatively the title compound as a yellow solid. LC-MS (ESI, m/z): 291 $[M+H]^+$.

**Scheme S3. Synthesis of ALG-097161[a]**

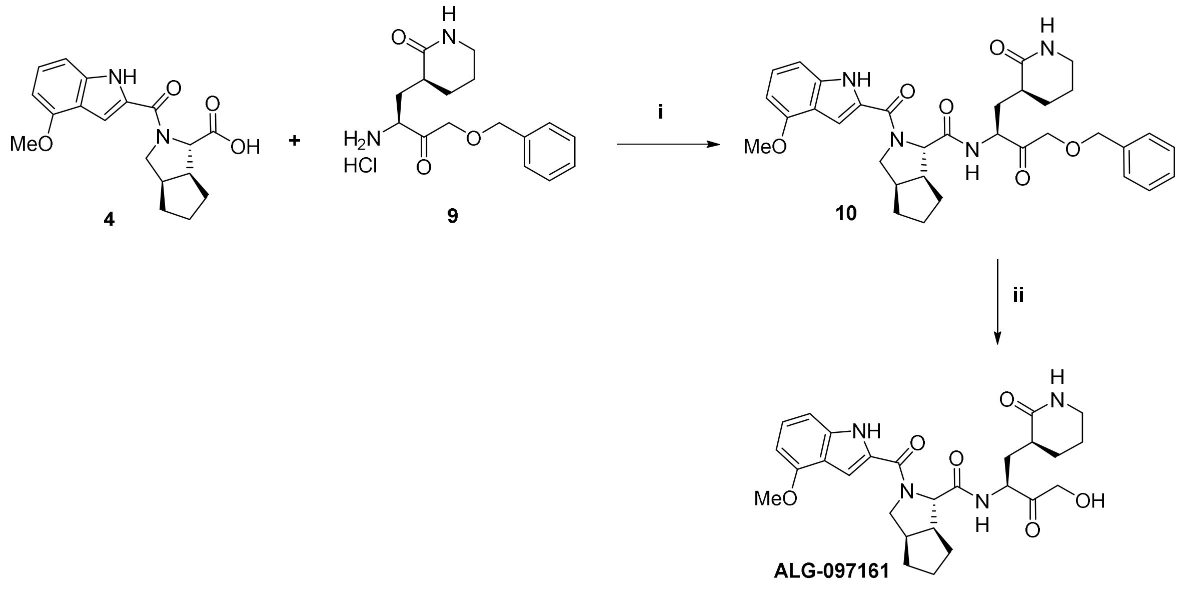

[a]Reagents and conditions: (i) TCFH, NMI, $CH_3CN$, 18%. (ii) $H_2$, Pd/C, EtOH, 14%.

**(1*S*,3a*R*,6a*S*)-*N*-((*S*)-4-(Benzyloxy)-3-oxo-1-((*S*)-2-oxopiperidin-3-yl)butan-2-yl)-2-(4-methoxy-1*H*-indole-2-carbonyl)octahydrocyclopenta[*c*]pyrrole-1-carboxamide (10)**

To a mixture of **4** (250 mg, 0.861 mmol), **9** (250 mg, 0.861 mmol) and chloro-*N*,*N*,*N'*,*N'*-tetramethylformamidinium hexafluorophosphate (265 mg, 0.947 mmol) in acetonitrile (6 mL) cooled at 0°C was added NMI (353 mg, 4.31 mmol). The reaction mixture was stirred at rt for 1 h. The reaction mixture was diluted with water (3 mL) and extracted with EtOAc (3 x 20 mL). The organic phases were combined, washed with brine (2 x 20 mL), dried over $Na_2SO_4$, filtered and concentrated under reduced pressure. The residue was purified by preparative TLC using 5% MeOH in $CH_2Cl_2$ as eluent to afford 95 mg (18%) of the title compound as a yellow oil. LCMS (ESI, m/z): 601 $[M+H]^+$.

**(1*S*,3a*R*,6a*S*)-*N*-((*S*)-4-Hydroxy-3-oxo-1-((*S*)-2-oxopiperidin-3-yl)butan-2-yl)-2-(4-methoxy-1*H*-indole-2-carbonyl)octahydrocyclopenta[*c*]pyrrole-1-carboxamide (ALG-097161)**

To a solution of **10** (90 mg, 0.150 mmol) in EtOH (5 mL) was added Pd/C (90 mg). The reaction mixture was stirred at rt overnight under $H_2$ atmosphere. The reaction mixture was filtered through a celite pad and the solids were washed with EtOH (20 mL). The filtrate was concentrated under reduced pressure. The residue was purified by preparative HPLC to afford 11 mg (14%) of the title compound as a white solid. LCMS (ESI, m/z): 511 $[M+H]^+$. $^1$H NMR (400 MHz, DMSO-*d*$_6$, 80 °C) δ 11.21 (s, 1H), 8.38 (s, 1H), 7.04 - 7.13 (m, 3H), 6.87 (s, 1H), 6.49 - 6.51 (d, *J* = 7.2 Hz, 1H), 4.46 - 4.69 (m, 3H), 4.17 - 4.20 (m, 2H), 3.95-

4.15 (m, 1H), 3.87 (s, 3H), 3.69 - 3.73 (m, 1H), 2.90-3.20 (m, 2H), 2.65 - 2.70 (m, 1H), 2.70 - 2.80 (m, 1H), 2.25 - 2.26 (m, 1H), 2.09 - 2.12 (m, 1H), 1.48 - 1.95 (m, 11H).
